# The 3D micro- and nanostructure of the apical extracellular matrix in *Drosophila*

**DOI:** 10.64898/2026.07.27.740909

**Authors:** Talha Alkhateeb, Jamal Ibrahim, Vincent Richter, Andreas S Thum, Matthias Behr

## Abstract

The apical extracellular matrix (aECM) serves as a dynamic scaffold, orchestrating tissue growth, cellular communication, and wound repair, and thus functions as a powerful biomaterial. In this study, we explored the intricate three-dimensional landscape of the *Drosophila* larval aECM using advanced 3D Volume electron microscopy. This technique opens a window into understanding and mimicking the diversity of aECM surfaces and architectures of the exoskeletal cuticles. We reconstructed the striking forms of denticles, hairs, and sensory organs, including double sensilla and Keil’s organs, at the epidermal surface. Our reconstructions brought the larval tracheal system and the sponge-like felt chamber (Filzkörper) of the spiracles into sharp relief, revealing a labyrinth of tiny cavities surrounding a central tube that extends from the posterior opening to the tracheal lumen. When examining the ultrastructure of transport-active anal pad cells, we discovered an inverted pattern of cuticular pore-canal structures, which supports their suggested role in osmoregulation across the aECM. Our findings further suggest that ordinary epidermal cells weave an intricate network of pore canals, enabling the orchestration of material distribution throughout the entire aECM. Moreover, we reconstructed tendon cells and chordotonal organs that form a branching network of numerous filamentous intracuticular fibers that weave through the aECM, linking the exoskeleton to muscles and sensory organs. By thoroughly mapping the intricate ultrastructural features of the aECM, this study opens new avenues for exploring how its protective, mechanical, and sensory roles intertwine, inspiring the next generation of tailor-made biomimetic materials.

## Introduction

The extracellular matrix (ECM) covers the basal and apical surfaces of epithelia in animals. ECMs are molecularly and structurally diverse, ranging from thin glycocalyx layers to thick exoskeletons, but all provide mechanical support, shape tissues, and protect against environmental factors ^1–4^. In vertebrates, the apical ECM (aECM) appears as a glycocalyx, mucin layer, or gel-like matrix in organs such as the respiratory tract, intestine, and renal tubules. Here, it functions as a diffusion and permeability barrier, supports signaling proteins, and buffers against shear forces ^2^. In invertebrates, the aECM forms an outer shell, such as the collagen- and chitin-rich cuticles of nematodes and arthropods ^5,6^. These layers serve as the first line of defense against mechanical stress, pathogens, dehydration, and chemical exposure, while also shaping sensory and communicative surfaces ^4,7,8^. Specialized aECM layers also develop transiently during embryogenesis and molting, contributing to the formation of tubes and intestine ^2,8,9^. In both vertebrates and invertebrates, the aECM connects to the cytoskeleton through transmembrane proteins and cell junctions, transmitting mechanical forces, distributing tissue tension, stabilizing epithelial integrity, and sensing environmental cues ^11^. Further, aECM regulates the degradation and regeneration of morphogenetic processes, including elongation, curvature, and expansion of epithelial organs ^12–14^.

The chitin-based aECM cuticle (exoskeleton) of arthropods is a natural composite of biopolymers and minerals from diverse materials, architectural styles, and properties ^6,15^. It is multifunctional, serving as both a protective barrier and an exoskeleton equipped with various tools and sensors ^16–18^. *Drosophila melanogaster is a* well-established genetic model for studying epithelial aECM cuticle ^1,3,7^. The outermost envelope layer contains protein-bound lipids, including cuticular hydrocarbons (CHCs), while the underlying epicuticle layer consists of proteins and lipids. Together, both participate in waterproofing and barrier functions ^19,20^.

The innermost and thickest layer, the chitinous procuticle, interfaces with living cells and anchors the cuticle at the apical cell membrane, enabling the exoskeleton’s function ^13,16,17,21,22^. Effective structural and functional organization depends on strong interconnection among all three layers ^23^. The epicuticle supports procuticle arrangement and cuticle stiffness, while the delivery of substances to the outer two layers requires an intact procuticle and pore canals ^19,23,24^. On the molecular level, numerous cuticular proteins and enzymes enter, upon secretion, the apical ECM to organize the chitinous aECM that constitutes non-lamellar and lamellar chitin/chitosan-protein matrices ^1,3,25^. This structural organization occurs at the cuticle assembly zone (deposition zone) and involves the synthesis and self-assembly of chitin chains into fibers, followed by protein-mediated polymerization into specific molecular structures that contribute to higher-order architectures ^10,16,23,25,26^. Disorganization of the aECM architecture leads to microcracks under mechanical pressure, exoskeleton detachment, insect slackness, vulnerability to mechanical, toxic, and pathogenic stress, desiccation, and ultimately rapid lethality ^17,19,27,28^. This makes the aECM a promising field for next-generation insecticides and bioinspired materials research ^15,29^.

The *Drosophila* respiratory system possesses non-lamellar chitinous aECM, offering greater flexibility. Tracheal cells secrete their aECM during late embryogenesis towards the apical lumen, where a thin chitinous cuticle forms an air-resistant, water-repellent barrier, enabling gas flow within the tube lumina ^1,30,31^. The tracheal airways require cuticular taenidial folds that run perpendicular to the lumen length, preventing collapse ^31,32^. In first-instar larvae, the two posterior spiracles (psp) are located on abdominal segment A8 at the posterior end of the larva. The spiracular opening, the stigma, is surrounded by four spiracular hairs, each composed of two neural and two support cells, consistent with the structure of a type I external sensory organ ^18^. The spiracles serve as primary respiratory openings, facilitating gas exchange ^31–33^. The air flows into the spiracular chamber, which contains a highly refractile filter called the filzkörper, which is lined by a bed of fine cuticular threads (felt) secreted by spiracular chamber cells that connect to the main tracheal tubes (dorsal trunk). This filzkörper meshwork of hydrophobic chitinous cuticle acts as an air filter that avoids liquid and pathogen entry, as well as dehydration ^32,34–36^. Spiracles are associated with three unicellular glands of elevated transcriptional and secretory activity. The ultrastructural features of the spiracular glands are consistent with a role in lipid secretion. This secretion is thought to confer hydrophobicity to the spiracle surface and to trap small particulate matter, helping protect the tracheal opening from water and debris ^37^. Although the morphologies of the trachea and spiracles have been studied in embryos, neither the 3D ultrastructural architecture of the spiracular (felt) chamber nor the reconstruction of the tracheal cuticle of *Drosophila* first-instar larvae is known.

The epidermal aECM contains a twisted plywood structure (Boulidgard structure) of chitin fibers with continuous rotation in the lamellar sheets ^16,38^. The orientation of those fibers affects the physical properties of the aECM ^15^. Within the aECM pore canals traverse the chitinous lamellae, and their structure likely depends on the orientation of the chitin fibers ^23,39^. Although their function is still debated, they have been shown to transport lipids and proteins to the outer layer ^40^, likely detect cracks in the cuticle for wound response ^41^, and may contribute to mechanical reinforcement ^42^. Previous studies of pore canal suggest a model in which lipid-transporting pore canals originate in the cytoplasm, cross the apical membrane, and eventually terminate at the surface of the envelope they perforate ^43^. In contrast, chitin-containing pore canals extend from the plasma membrane to the epicuticle region ^44^. However, their 3D structural formation within the aECM remains poorly understood.

Epidermal cells of *Drosophila* are intrinsically mechanosensitive and modulate nociceptive behavioral responses. Numerous Somatosensory neurons (SSNs), including mechanosensory (C3da) and (C2da) neurons, and proprioceptive (C1da) neurons, innervate the epidermis, in addition to (C4da) nociceptors, while chordotonal neurons are attached to the epidermis ^45,46^. Thus, two main categories of sensilla form at the epidermis: chordotonal organs (stretch receptors) and external sensilla (mechanoreceptors and chemoreceptors). The accessory cells in the external sensilla remain in the epidermis, forming a cuticular apparatus that receives stimulation. The chordotonal organs are primarily proprioceptors that remain subepidermal, as they detect internal mechanical stimuli related to body position and movement. While they are predominantly proprioceptive, they can also detect a wide range of external stimuli^47–49^. In *Drosophila* larvae, mono- and pentascolopidial adhere to the lateral body and provide touch sensitivity and sensory feedback to the locomotor circuit during locomotion ^18,50^.

The functional unit of a chordotonal organs consists of four cells: a neuron, a scolopale cell, a cap cell, and a ligament cell. At the core of each scolopidium, a bipolar neuron extends its dendrite into the scolopale cell. In clustered organs such as the larval lateral pentascolopidial organ (lch5), each of the five neurons features a teardrop-shaped cell body and a single dendrite oriented dorsally and posteriorly. The scolopale cell, a glial-type accessory cell, ensheathes the neuron’s dendrite and, together with the neuron, forms the organ’s structural and functional core. The cap cell is located on the dorsal side of the scolopidium and distally connects to the scolopale-neuron unit, anchoring it to the epidermis via an epidermal cap attachment cell. The ligament cell is positioned ventrally and extends the organ toward its opposite anchor point in the cuticle, mechanically linking the scolopidium to the body wall and allowing body movements to stretch the sensory unit ^51^. Proper function of chordotonal organs as stretch receptors depends on their stable anchoring ^52^, a process that remains poorly understood.

Locomotion depends on the synergy of soft and hard tissues ^53^. Vertebrates connect muscles to the endoskeleton via tendons. In contrast, invertebrates, particularly arthropods, use specialized muscle-tendon connections to securely anchor muscles to their aECM (exoskeleton), providing both movement and structural stability. In the absence of an internal skeleton, specialized epidermal cells, called tendon cells (tenocytes, apodemes), act as intermediaries, connecting muscles basally and the chitinous exoskeleton apically ^21,54,55^. At the tendon-cell junction, specific intercellular structures, such as myotendinous junctions (MTJs) and apical hemi-adherens junctions (aHAJs), provide physical links between muscles and the tendon exoskeleton, respectively, thereby ensuring force transmission during locomotion ^54^. Apically, tendon cells differ from neighboring epidermal cells; they secrete electron-dense protrusions, called intracuticular fibers (ICF, tonofilaments, tonofibrillae), directed from the apical cell membrane into the cuticle ^54,55^. Recently, we demonstrated that the loss or disturbance of ICFs results in detachment of the chitinous aECM from epidermal cells, leading to severe locomotor deficiencies in *Drosophila* embryos and larvae ^17^. Thus, detailed knowledge of aECM 3D structure offers insights into future biomimetic material design, revealing how material compositions anchor to withstand rising mechanical forces. Yet a detailed structural analysis of these crucial elements and their connections to the aECM is missing.

By fusing high-resolution 3D Volume structural analysis with the power of AI-driven modeling, our study uncovers the ingenious strategies nature uses to shape aECM, revealing a stunning array of forms and functions. We offer a sweeping exploration of cuticle architectures and the sensory neurons associated with them. We further delve into the unique appearances and roles of epidermal, tracheal, and spiracular cuticles, guided by their internal morphology and nanostructures. We also highlight the intricate ramifications and connections within internal cuticle structures, including pore canals and intracuticular fibers at the epidermis, anal pads, tendon, and sensory cells. These vivid reconstructions not only deepen our understanding of evolutionary biology and ecology but also open new avenues for biomimicry. Our discoveries may lay the groundwork for bioinspired innovations in micro air vehicles, advanced fiber-reinforced materials, water-repellent surfaces, and robust protective coatings.

## Results

Microstructural characterization of tissues has relied heavily on high-resolution images produced by electron microscopy. However, the full larval body volume we used was established by the ssTEM technique (serial sections imaged with a TEM in scanning mode), which was enabled by an automated ultramicrotome ^56^ providing a precisely measured whole-animal volume. The first use of this technique has advanced our understanding of the complete enteric nervous system and the larval sense organs of *Drosophila* larvae ^18,57^. We used this first-instar ssTEM dataset of 14,448 images (∼3.5 TB; resolution: 99600x31600 pixels) for subsequent AI-based aECM reconstructions using the Dragonfly 3D World software (2024.1).

To initially handle the large volume of data, the image series was divided into three data blocks (anterior, median, and posterior). In addition, to process the large number of images of each data block, the pixel resolution was reduced by a factor of 10. Finally, all images from the three blocks were stitched together, and this volume was used for subsequent analysis.

After training the deep learning model to segment the epidermal cuticle and trachea, Dragonfly provided a results diagram showing the model’s training and validation losses over several epochs (Fig. S1A). Initially, the loss values are high, indicating an inadequate model fit. Over time, a clear, continuous decline in both curves can be observed. After the first ten epochs, there is a substantial decrease in losses, with the training loss decreasing more uniformly than the validation loss. At the end of training, both curves reach values below 0.01, indicating an acceptably low error rate. These results demonstrate the model’s capacity to reliably segment both the epidermal cuticle and the tracheal lumen, making it suitable for quantitative analysis of nanostructural data. We applied the deep learning model to the image data to label the epidermal cuticle. The labeling was highly precise and reliable, enabling detailed morphological analysis (cuticle marked in blue; Fig. 1A-C; Fig.S1B). We used the same AI model to identify the tracheal tubular structures. First, we marked the tracheal lumen and trained the model to recognize it. In the end, the AI model successfully and accurately recognized the entire tracheal lumen (marked in yellow; Fig. 1D). These results confirm the model’s high performance in segmenting complex ultrastructural data. We then used the deep learning tool to create 3D models of the epidermal cuticle and tracheal lumen from the AI-marked areas. For optimal visualization, we rendered the models, adding lighting and shadows to make the morphological structures more visible.

**Figure 1.**
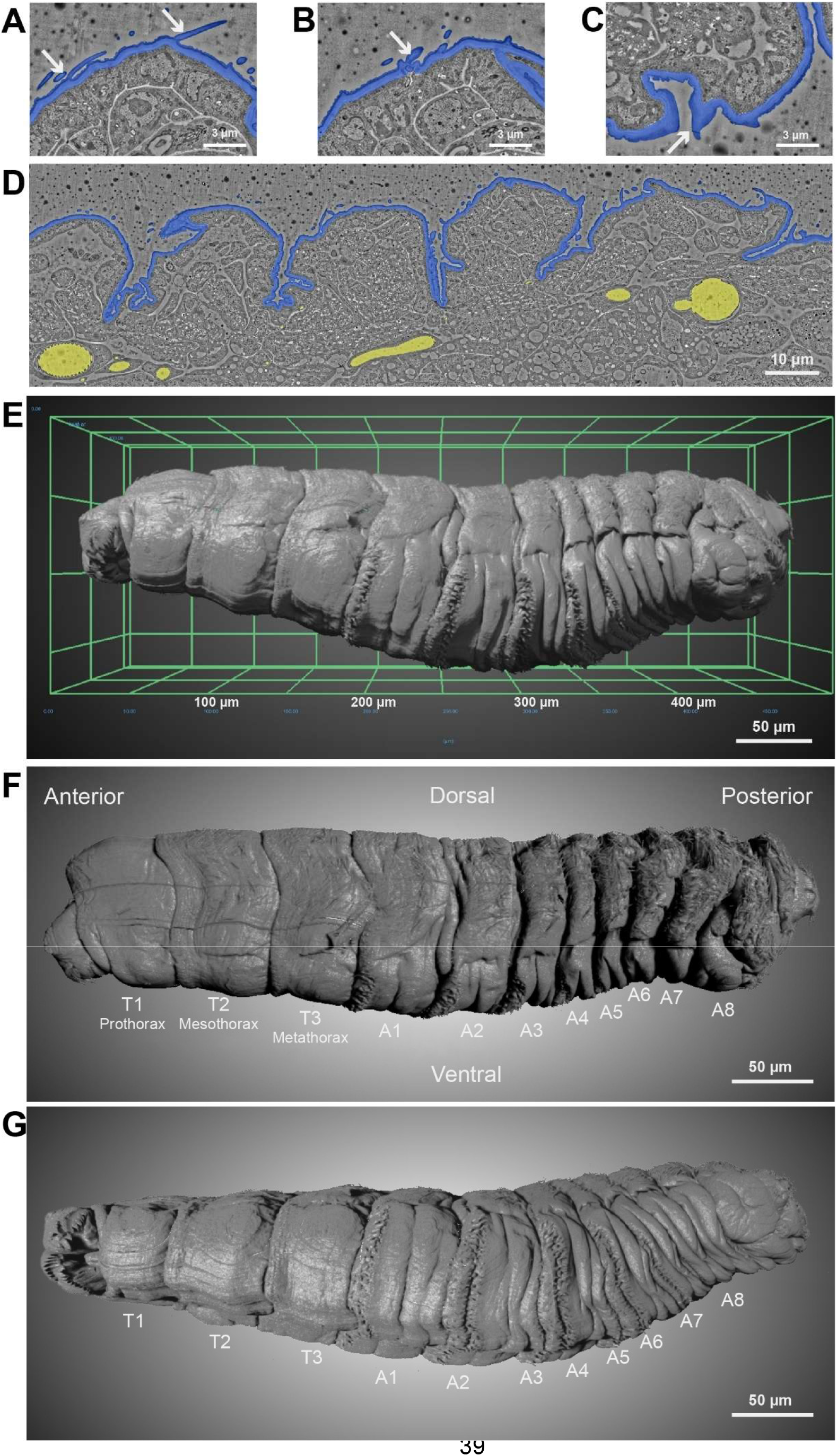
(A–D) Electron micrographs of the larval cuticle segmented by deep learning to identify hair-like structures, sensilla, denticles, and tracheal lumens. Cuticle segments appear in blue, and the tracheal lumen in yellow. **(E)** Ventrolateral view of the 3D larval model after cuticle rendering, with a micrometer scale bar. **(F, G)** 3D lateral view of the L1 larva. The pseudocephalon is visible at the anterior (left), and the posterior spiracles at the posterior (right). Rows of denticles in abdominal segments (A1–A4), finer thoracic denticles, and dense hairiness in segments A2–A8 are visible. Scale bars are included.

Rendering the dataset reveals 3D reconstructions of the segmental larval body organization and characteristic cuticle surface structures. A 3D lateral view of the first instar shows the body segments (anterior left; Fig. 1E-G). The rows of denticles are clearly visible in the abdominal segments (A1–A4). In the thoracic segments (T1-T3), the rows of denticles appear thinner, and the individual denticles are smaller. Hairs are particularly dense in the dorsal region of the larva, especially in segments A2–A8 and around the posterior spiracles at A8. The pseudocephalon is visible at the anterior end ^33^.

Larvae sense their environment through specialized external organs that detect specific sensory cues, including olfactory, gustatory, temperature, and mechanosensory signals ^58–60^. To illustrate larval morphology in detail, we reconstructed various characteristic structures of the epidermal cuticle in the first instar (Fig 2). Central morphological features crucial for the functionality and behavior of the larva became visible, such as the cirri and mouth hooks of the pseudocephalon (Fig. 2A). The cirri consist of sensory bristles that contribute to tactile perception and food intake ^45,61^. The mouth hooks are chitinized structures that mechanically break down food ^62^. The Keilin’s organ is visible in the ventral thoracic segments T1 to T3 (Fig. 2A). This mechanosensory organ plays a central role in tactile perception during larval development in insects. It typically consists of a group of three to five sensory cells that respond to tactile stimuli ^6^.

**Figure 2.**
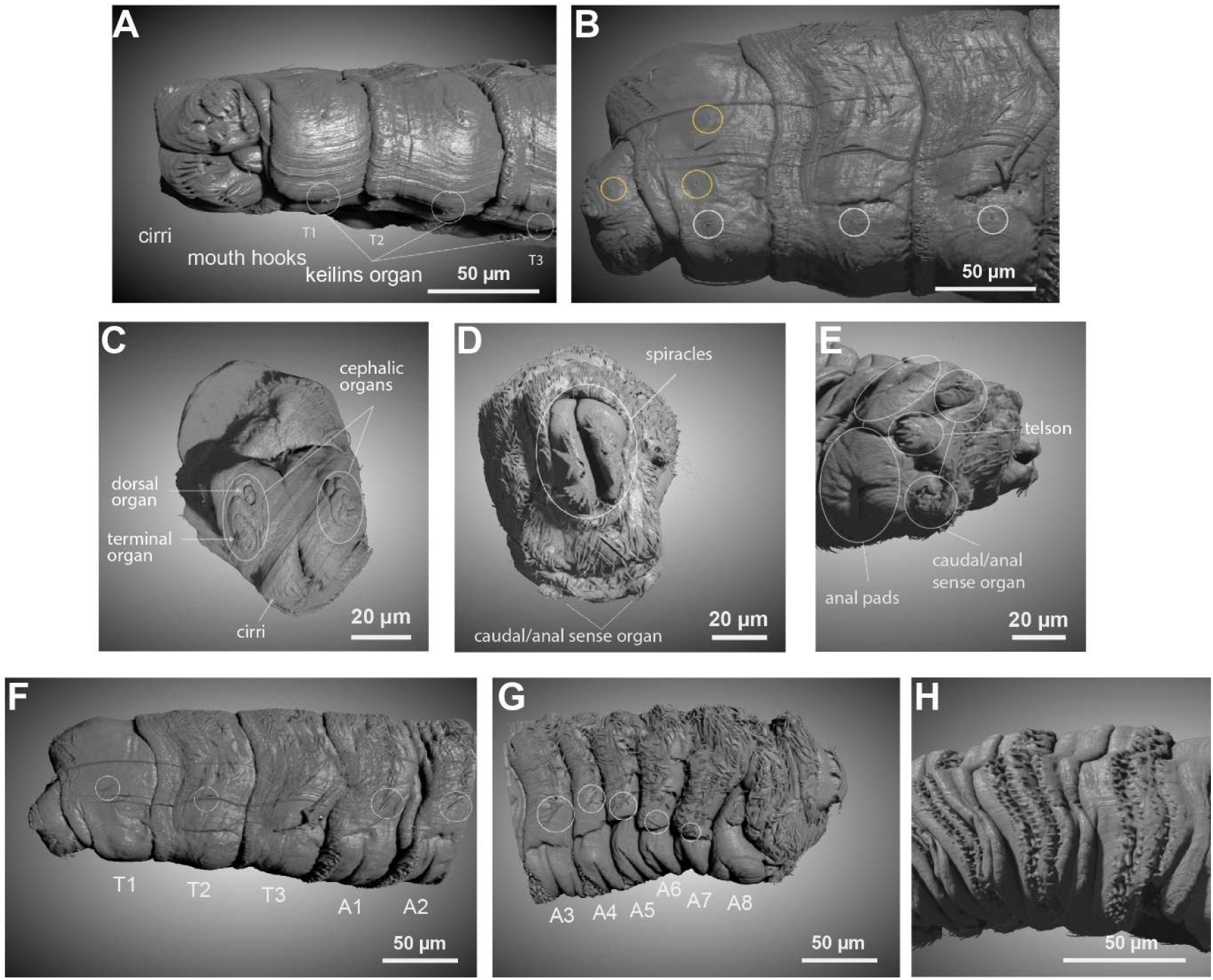
(**A**) The cuticle rendering presents an anterolateral view of the larval head, highlighting the cirri and mouth hooks. **(B)** The anterolateral view displays three knob sensilla (circled in white) and three papillae sensilla (circled in yellow). **(C)** The morphology of the larval head is shown, with the anterior side and sensory organs visible. The dorsal side is at the top and the ventral side at the bottom. Cirri surround the mouth opening. **(D)** The terminal segments of first-stage larvae are depicted. Two spiracles of the tracheal system protrude from the body, and the caudal or anal sensory organ is visible ventrally. **(E)** A posterior-lateral view of the anal sensory organs, caudal organs and telson and anal pads (all circled). **(F, G)** Hair sensillae are clearly visible in thoracic segments T1 and T2. In segment T3, they are not apparent due to cuticle damage. Hair sensillae are also present in abdominal segments A1 to A7 (circeld). **(H)** The ventral view of the epidermal denticles is shown. Scale bars are included.

### Reconstructions and 3D projections of larval sensilla at the cuticle

The 3D reconstruction of the cuticle of the anterior region of the larva shows several sensory organs, including terminal and dorsal organs, the main cephalic organs of the larvae (Fig. 2C). These organs harbor mechanoreceptors, chemoreceptors, and thermoreceptors that are essential for perceiving environmental stimuli. They allow the larva to respond appropriately to these sensory modalities, orienting itself in its environment ^45,64,65^. The cuticle reconstruction of the posterior end shows segments A9–A11, the anal pads, and the telson. where caudal/anal and telson sensory organs are recognizable (Fig. 2D,E). These sensory organs respond to environmental stimuli and assist the larva in orientation ^33^. In addition to sensory organs, body hairs and denticle bands are particularly prominent in the reconstruction (Figures 1F,G; 2H).

Thoracic and abdominal sensilla form during *Drosophila* embryogenesis. This includes accessory cells at the apical portions of sensilla, which form the cuticle apparatus for receiving sensory stimuli during larval stages ^63^. Hair sensilla are visible in the thoracic and abdominal segments (Fig. 2B, F, G). The hair sensilla play a central role in the perception of mechanical and chemical stimuli, enabling the larva to respond specifically to its environment and support processes such as orientation, locomotion, and foraging ^18,66^.

The cuticle of insects is further decorated with non-sensory hairs (microtrichia or trichomes) (Figures 1F,G; 2G). Trichomes enhance the cuticle’s hydrophobicity, which aids in larval locomotion to reduce or increase surface friction. The denticle bands are chitinous structures that serve to propel the larva on solid surfaces and are characteristic of each segment (Fig. 2H).

To detect the 3D ultrastructure of sensilla cuticle, we trained a new AI model to generate high-resolution 3D models of specific epidermal cuticle structures, which we subsequently rendered. This revealed three types of sensory organs located on the thoracic and abdominal segments of the larva: hair sensilla (including single-and double-hair sensilla), knob sensilla, and papilla sensilla (Fig. 3A-D). These mechanosensory sensilla occur both individually and in more complex sensory organs ^18,59,63^. The 3D projections of the ultrastructure images and sections reveal that the sensilla are lined by the cuticle (Fig. 3E-L). A more detailed analysis of the ultrastructure of 3D projections shows that hair sensilla are sourrounded by the three cuticle layers, including the chitinous procuticle, which appears to extends in thickness adjacent to the tubular body (Fig. 3H,I,L). The tubular body of the papilla and knob sensilla passes the procuticle and reaches into the epicuticle (Fig. 3F,G,J,K). Thus, the diverse morphological structures indicate that sensory and mechanical elements are closely interconnected throughout larval development.

**Figure 3.**
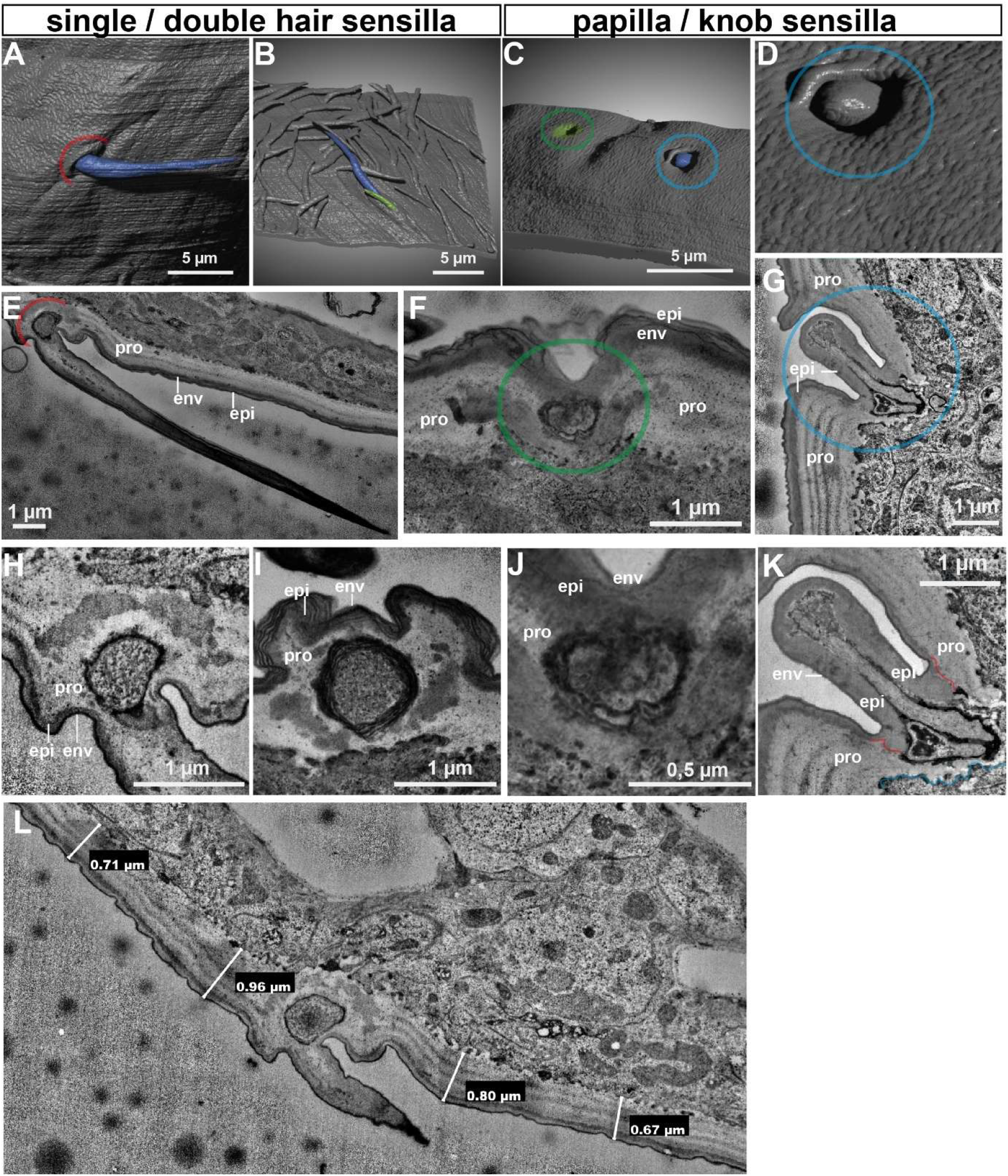
3D reconstructions and projections of the cuticle sensillae. **(A)** Hair sensillae in the first thoracic segment are stained blue. **(B)** Double hair sensillae in the first abdominal segment include a short, green-stained hair and a long, blue-stained hair. **(C,D)** Papillae sensillae (green) and knob sensillae (blue) are located in the anterior region of the pseudocephalon. **(E-L)** 3D projections and magnifications of the cuticle at the hair (E,H,L), double hair (I), papilla (F, J), and knob sensilla (G,K) are shown. Scale bars are included.

### Reconstruction of larval tracheal tubes

The tracheal tubes are arranged in a stereotypical, segmental pattern and branch throughout the larval body that enables air transport to target organs ^31,32^. After AI training, Dragonfly generated a 3D model of the tracheal branch lumina (Fig. 4A). The program automatically transfers size measurements from TEM images to the corresponding section of the 3D model. To verify accuracy, distances between two points of the marked tracheal lumen were measured horizontally and vertically in a 2D section. The 3D model measurements matched those from the TEM images in all comparisons (Fig. S2). A detailed 3D reconstruction depicts the two main dorsal trunks of both hemispheres, which extend from posterior to anterior and terminate at the spiracles, and also visualizes the primary tracheal branches. Combined with the epidermal cuticle, the model highlights the close proximity of most tracheal branches to the epidermis as they spread along it (Fig. 4A, Fig. S3). Videos of the 2D stacks and 3D reconstructions highlight tracheal ramifications of ganglionic branches (Video 1-3). These 3D models provide a foundation for further quantitative analysis of the larval respiratory system. Additional FIB-SEM data of the tracheal tube shows in great details the oscillating cuticular wall (taenidial folds) of the trachea (Video. 4) with spacing between ridges about 2.7-3.1 um (Fig. 4B). The maximum diameter of the cross-section of the tracheal tube (major axis) is 6.8 µm (std = 1.1 µm) and the minimum (minor axis) 4.6 µm (std. 0.6 µm), the oscillations are clearly shown by the change in both diameters (Fig. 4B, bottom)

**Figure 4.**
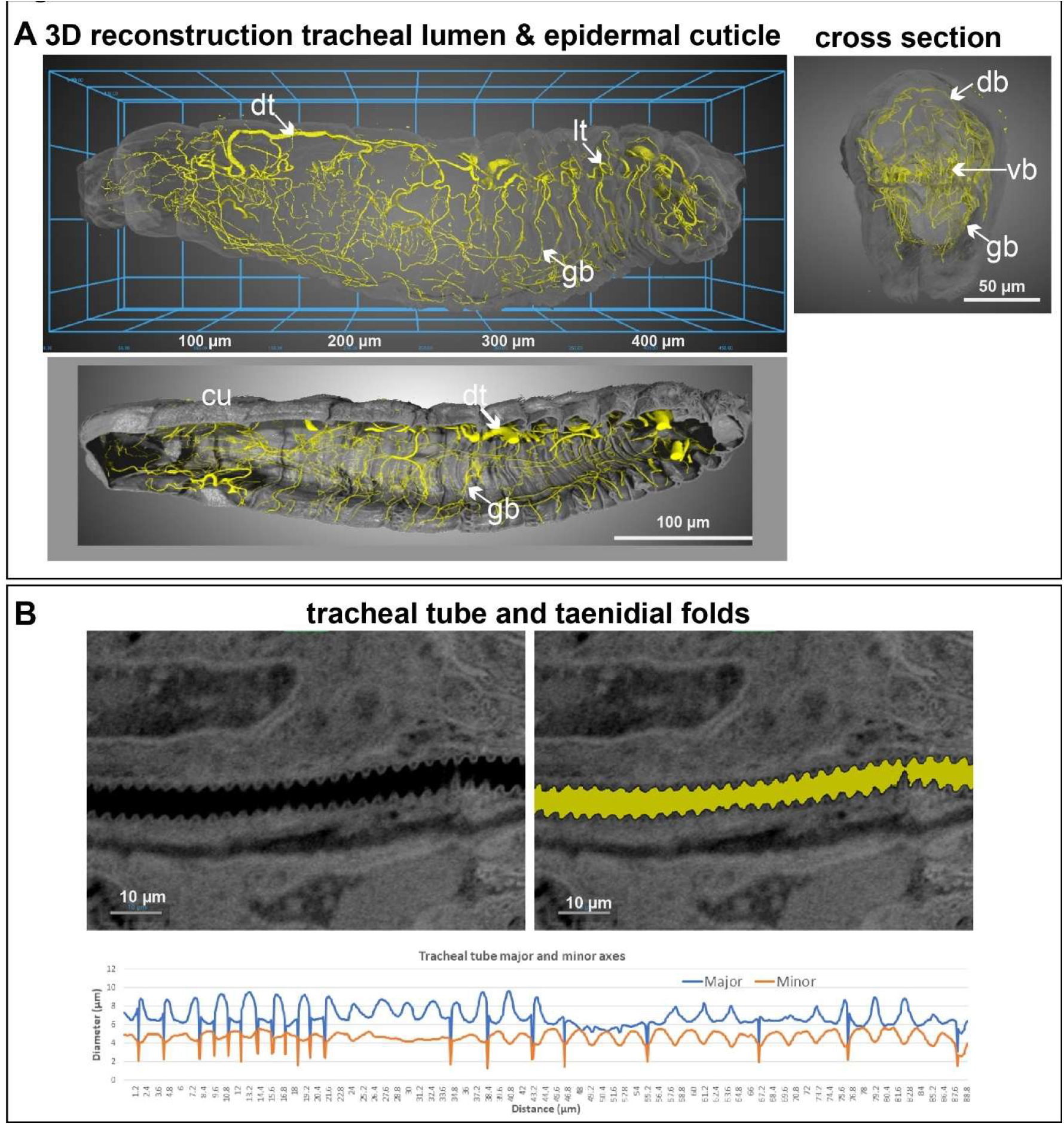
Morphology of the tracheal system. **(A)** Upper and lowar panel: lateral view of the rendered 3D tracheal system model (yellow) with a micrometer scale. The epidermal cuticle (cu) is marked in grey. dorsal trunk (dt), dorsal branches (db), ganglionic branches (gb), lateral trunk (lt), visceral branches (vb). Figures are oriented with the dorsal surface upward and the anterior to the left. The right images shows a reconstructions across the larval body. **(B)** FIB-SEM histogram section showing a tracheal tube. Left, backscattered image showing the tracheal tube after resampling the viewing plane. Right, the same slice in the left after segmenting the tracheal tube. Scale bar = 10µm. Bottom, plot showing the major and minor diameters of a tracheal tube measured along its full length. The graph shows regular, repeating fluctuations in both axes that reflect the ridges within the tracheal tube wall, the distance between these ridges is between 2.7 and 3.1 µm. Scale bars are shown.

### Reconstruction of larval posterior spiracles

The cuticle is a central element of posterior spiracles. The large spiracular chamber harbors cuticles that spread as thread- or hair-like structures ^32,37^ in the 2D images and connect to the dorsal trunk at each hemisphere (Fig. 5A). However, in three dimensions, the threads transform into a dense web of rod-shaped cuticle protrusions. In the 2D video of the spiracular chamber, one can trace how these protrusions extend from the apical side into the lumen (Video 5). Yet, it is the 3D reconstruction that reveals their complexity. Here, the protrusions not only extend into the lumen but also link with their neighbors, weaving a thin internal tube that runs from posterior to anterior through the entire spiracular chamber while creating a sponge-like, porous structure around it (Video 5; Fig. 5B-D). Cross-sections of the 3D reconstruction show that fused adjacent cuticle protrusions create small cavities around the central lumen (Fig. 5B). These cavities organize between the apical cell membrane and a central lumen and connect to the lumen through openings (Fig. 5E). This unique architecture forms a unit that resembles the texture of pipe cleaner felt (“Filzkörper”) inside the pipe.

**Figure 5.**
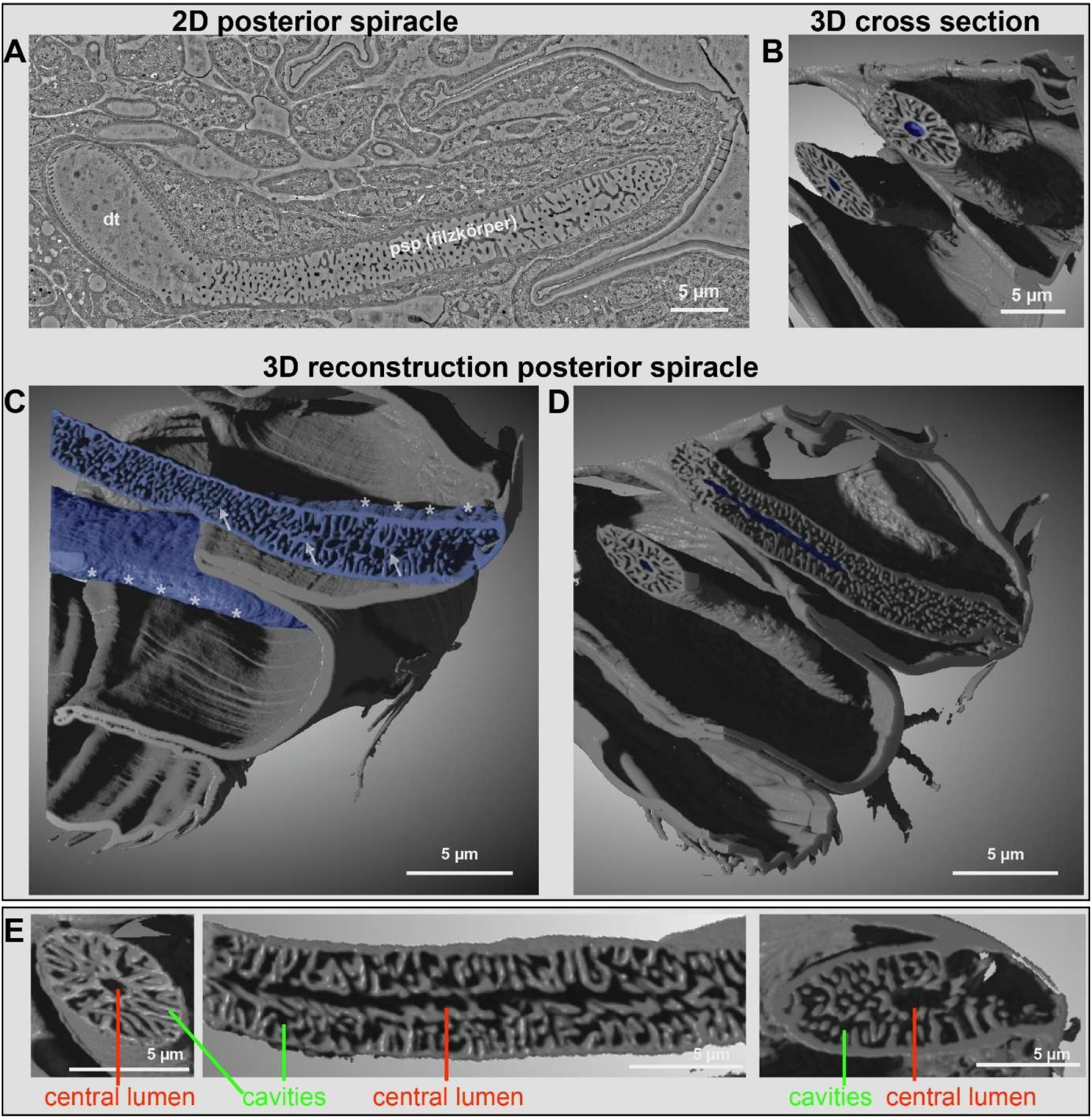
Reconstruction of the internal structure of the posterior spiracles. **(A)** 2D TEM image showing the posterior spiracle, the Filzkörper with cuticle threads, and the transition into the tracheal lumen of the dorsal trunk. **(B-E)** 3D reconstruction shows a cross-section of the spiracle chamber, highlighting the Filzkörper and the centrally positioned tube, both shown in blue. Two short video clips accompany this figure (B). The spiracle chamber and the Filzkörper are shown in blue. Stars indicate the spiracle chamber, while arrows indicate the Filzkörper (C). Illustration of the Filzkörper forming a tube that runs centrally through the spiracle chamber, shown in blue. Scale bars are indicated (D). Magnifications of reconstructions indicating the central lumen and the cavities of the Filzkörper (E). Scale bars are indicated.

### Reconstruction of larval anal pad cell and cuticle

Next, we reconstructed the cuticle of the anal pads, two symmetrical organs adjacent to the epidermis at the larval tail. The overall cuticle organization of the anal pads is specific. Anal pad cuticle contains lamellar chitinous procuticle and an outer envelope, but the epicuticle was not detectable when compared with neighboring epidermal cells (Fig. S4,S5). Also, anal pad cells are of specific appearance. The apical cell membranes form numerous closely spaced infoldings, many associated with mitochondria (S4). We reconstructed a single anal pad cell to understand more about the cell-specific distribution of mitochondria in relation to the apical cuticle (Fig. 6A). In 3D, mitochondria are fused into a large structure (Fig. 6A, light brown) distributed throughout the cytoplasm and extending to the apical membrane infoldings next to the cuticle. Ten additional smaller mitochondria are present at the apical cell membrane, too. Furthermore, electron-dense extracellular structures are located between the membrane infoldings, next to the mitochondria (Fig. 6A, black). In the 2D TEM images and 3D projections of the cuticle, we observed thick, potential electron-dense pore canal-like structures at the envelope. The 3D reconstructions show a high number of these structures protruding from the envelope into the procuticle (Fig. 6B-D). The thick canal-like structures protrude from the envelope into the procuticle and are connected with thinner canal-like structures that reach towards the apical cell membrane (Fig. 6D; Fig. S5). Thus, the canal-like structures connect the apical membrane with the outer cuticle surface, indicating the potential of dynamic transport activities across the cuticle.

**Figure 6.**
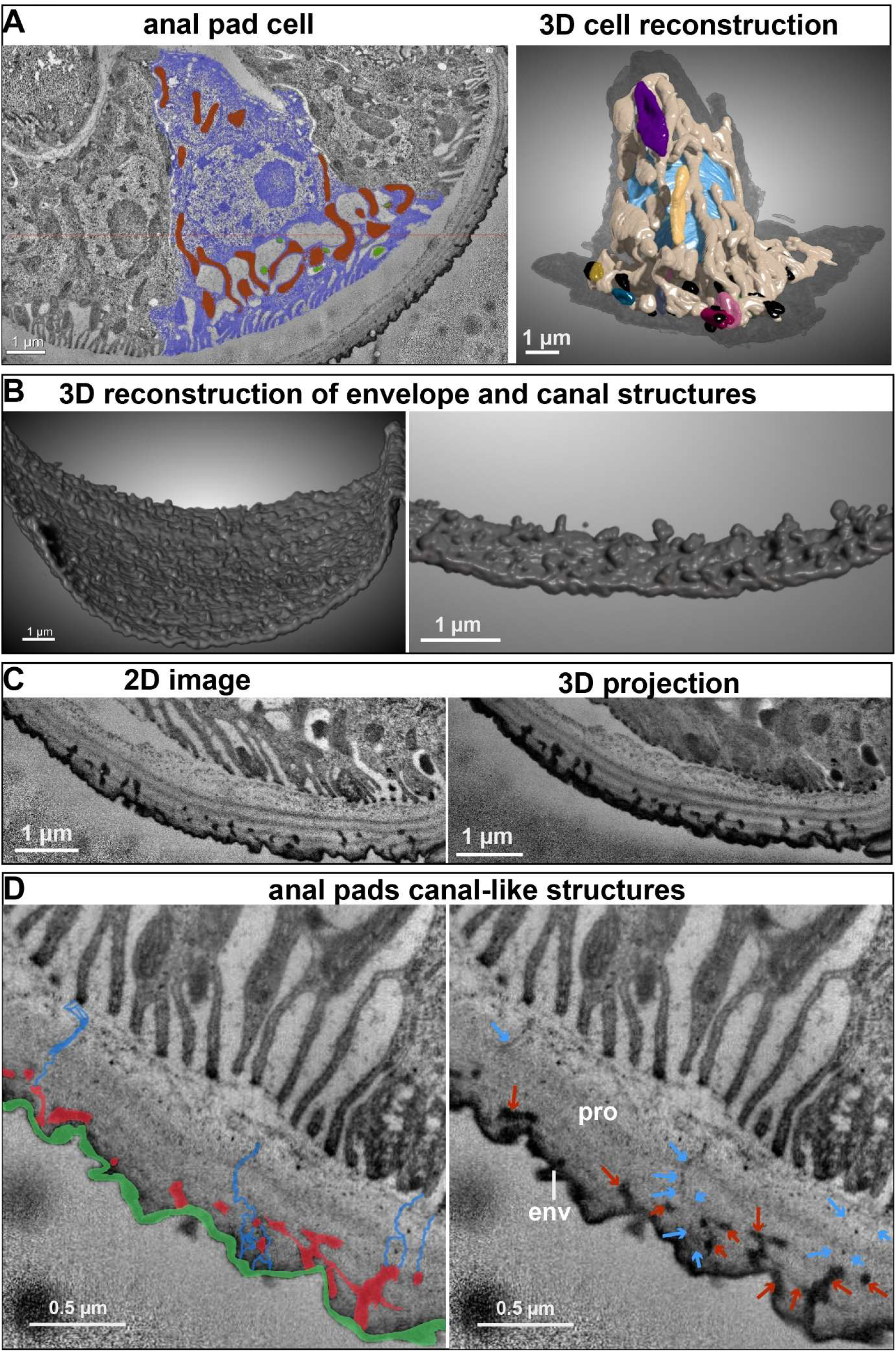
Reconstructions and projection of anal pad cell mitochondria and cuticle. **(A)** Anal pad cell is marked in blue and mitochondria in brown (2D TEM, left image). The 3D reconstruction reveals that mitochondria fused to a giant structure (light brown) within the cytoplasm reaching to the membrane infolding at the apical cell site (right image). Additional small mitochondria appear (purple, yellow, dark blue, red, pink, and gray). Nucleus is marked in blue, and dense structures (black) appear between membrane infoldings next to mitochondria. **(B)** A 3D reconstruction of the electron dense cuticle envelope shows rough surface appearance including likely many thick pore canal-like structures (left), with close-up on the right. **(C)** The cooresponding 2D TEM image and 3D projection of anal pad region of the reconstructions exhibits large electron dense canal-like structures reaching into the lamellar procuticle. **(D)** A close-up of TEM images of the anal pad cuticle reveals wide, interconnected pore canals marked in red (left and red arrows in the right image) and thinner canal-like structures (marked in blue in the left and blue arrows in the right image) that traverse the procuticle (pro). The similar unmarked pore canals are visible in Fig. S6. The envelope (env) is marked in green, while the underlying epicuticle is not detectable. Scale bars are indicated.

### Reconstruction of larval epidermal cuticular nanostructures

The epidermal cuticle also contains pore canal-like structures. We identified these structures in the *Drosophila* larva originating from the apical membrane, often at the undulae, extending through the layers of the cuticle to the envelope surface (Fig. 7A). The 2D TEM images show potential pore canal-like structures of different diameters. Central pore canal-like structures, with an approximate diameter of 20nm that, traverse the procuticle or run axially in the procuticle, connecting canals along the chitinous lamella (marked in blue; Fig. 7A). Additionally, numerous thinner canal-like structures, below 10 nm in diameter (marked in orange, Fig. 7A) connect to the central elements. Because 2D images can show only portions of these structures, we reconstructed them in a 3D model to assess their organization. The 3D view reveals a branched network of the central and thinner structures extending both horizontally and vertically within the cuticle, being connected to the apical membrane and the envelope (Fig. 7B; Video 6,7). Thus, thicker central structures in the procuticle run from the membrane across the chitin matrix lamellae, from which multiple thin structures are located in the procuticle and traverse the outer cuticle layers towards the surface (Fig. 7A,B; Fig. S6). We further confirmed that pore canal-like structures run through the cuticle with FIB-SEM (Fig. 8 and Video 8). Our data show that they can also be laterally oriented, i.e., running parallel to the cuticular wall, not only perpendicular (Video 8). In summary, the appearance of perpendicular and laterally canal-like structures indicates that the epidermis forms a transport system across the cuticle, suggesting that the cuticle is not rigid but can undergo dynamics that likely establish its formation, degradation, and preservation.

**Figure 7.**
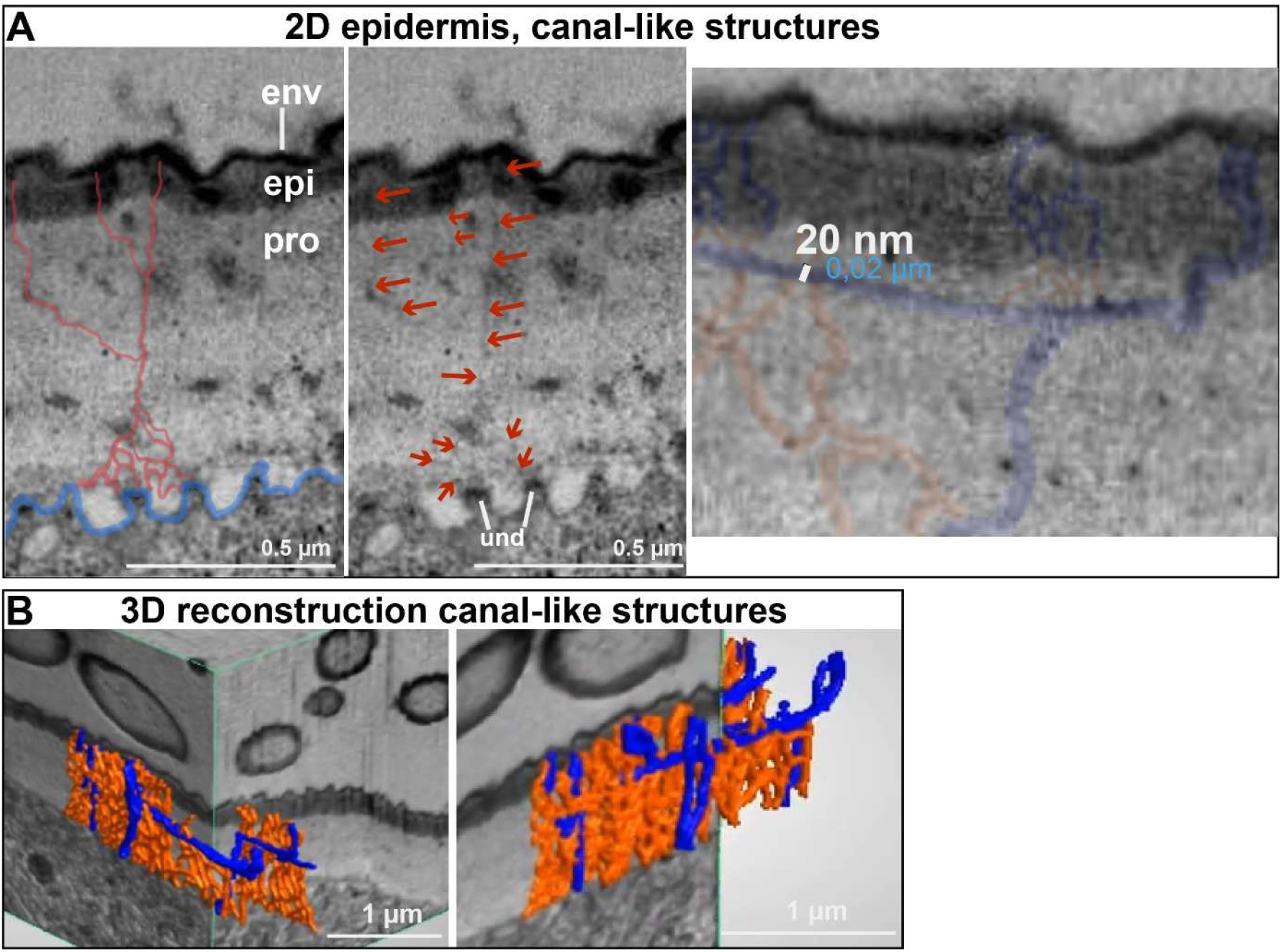
Pore canal-like structures in the epidermal cuticle. **(A)** TEM images display various pore canals in the epidermal cuticle, with thin canal-like structures (10nm) in red and thicker ones (20nm) in blue. Red arrows point to the corresponding thin canal-like structures (10nm). Additional unmarked images are provided in Figs. S6,S7. Thicker horizontal and vertical pore canal-like structures (right image, marked in blue) have a diameter of 20 nm. Right image: apical cell membrane is marked in blue and undulae (und) are indicated. **(B)** Both wider and thinner canals form a network as illustrated in the 3D reconsctructions. Thicker channels in blue and thinner ones in red. Two video clips accompany this figure. Scale bars are included.

**Figure 8.**
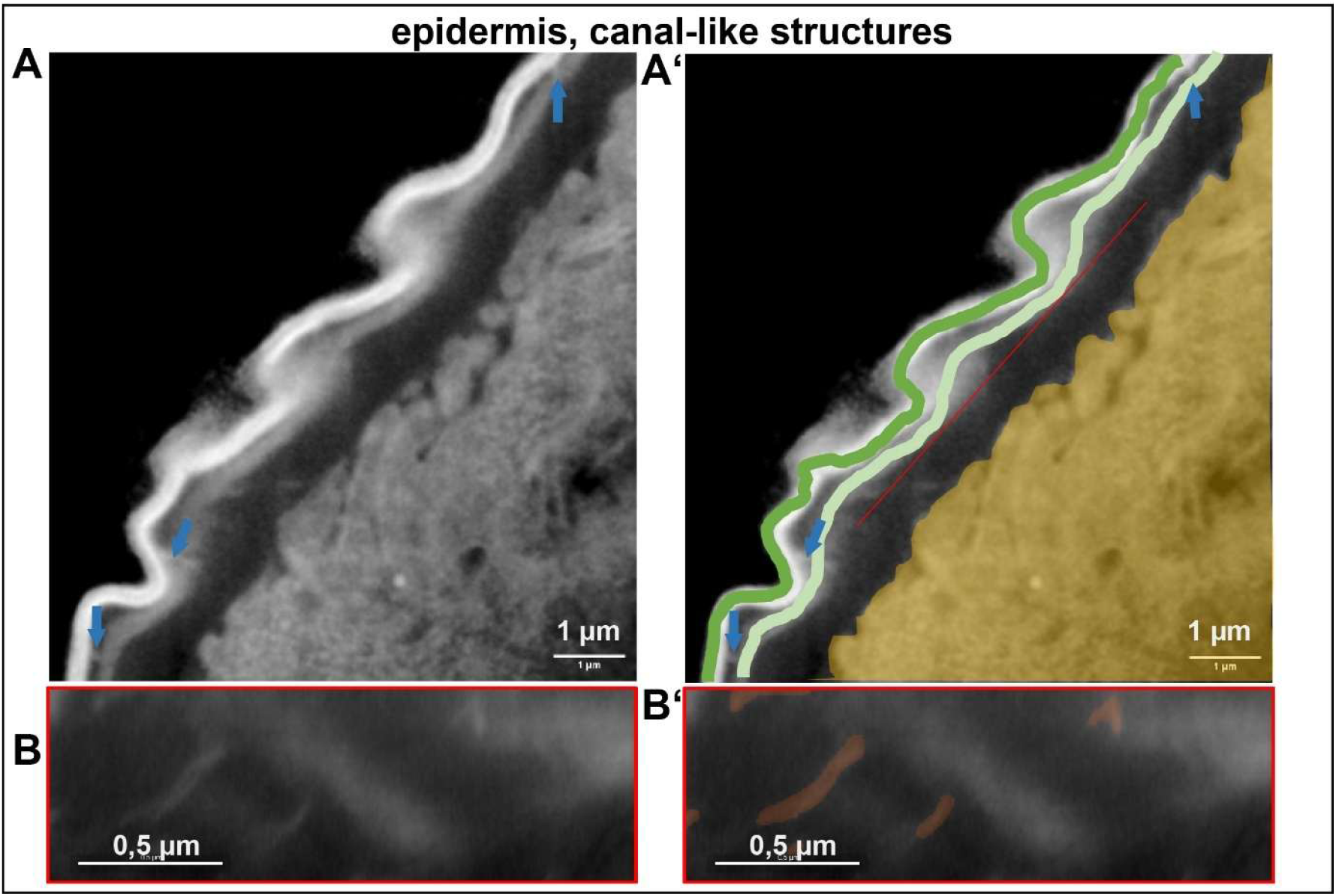
Epidermal cuticular pore canal-like structures. FIB-SEM slice showing to the outer cuticle in the 1^st^ install larvae. **(A)** Slice from backscattered detector showing the epithelium layer and the cuticle, **(A’)** the same slice in A highlighting the epithelium (orange) and the cuticle (green), arrows in both images point to the pores that traverse the cuticle through the assembly zone towards the outer cuticle, scale bar = 1 µm. **(B, B’)** The red line from image A’ points to the location from which image was resampled highlighting the pore canal-like structures, scale bar = 0.5 µm.

### Reconstruction of larval intracuticular fibers (ICFs) at tendon cells

The cuticle forms a resilient exoskeleton, tightly anchored to the muscle apparatus by specialized tendon cells at the epidermis. 2D TEM images, 3D projections, and 3D reconstructions revealed somatic muscles that attach the tendon cells (located at apodemes) basally via myotendinous junctions (Fig. S7A,B). The 2D TEM images, and 3D projections show that the tendon cells are connected apically with the cuticle via apical hemi-adherens junctions (aHAJs) and ICFs (Fig. 9A; Fig S7). These aHAJs and ICFs (intracuticular fibers; tonofilaments) play key roles in attaching the cuticle to the larval tendon cells for proper locomotion ^17,54^. The 3D reconstructions of these fibers in 2D TEM images exhibit an ICF network that connects the tendon cell to the exoskeleton. At each aHAJ attachment site, we observe large ICFs reaching into the cuticle, from which highly branched, numerous thin filamentous ICFs spread into the cuticle, suggesting an anchorage-like function (Fig. 9B; S7C). This filamentous ICF anchorage forms an intricate, branching network that weaves through all cuticle layers and reaches into the outermost envelope (Fig. 9B,C; Fig. S8C). Analogously, numerous filamentous-ICFs anchor the large ICF to the aHAJs of the apical cell membrane (Fig. 9C,D). Additionally, filamentous-ICFs anchor large ICFs to the apical cell membrane adjacent to aHAJ (Fig. 9B; S7C). Our data indicate that a branched ICF network provides the anchor for the tendon cell to the exoskeleton (Fig. 9E).

**Figure 9.**
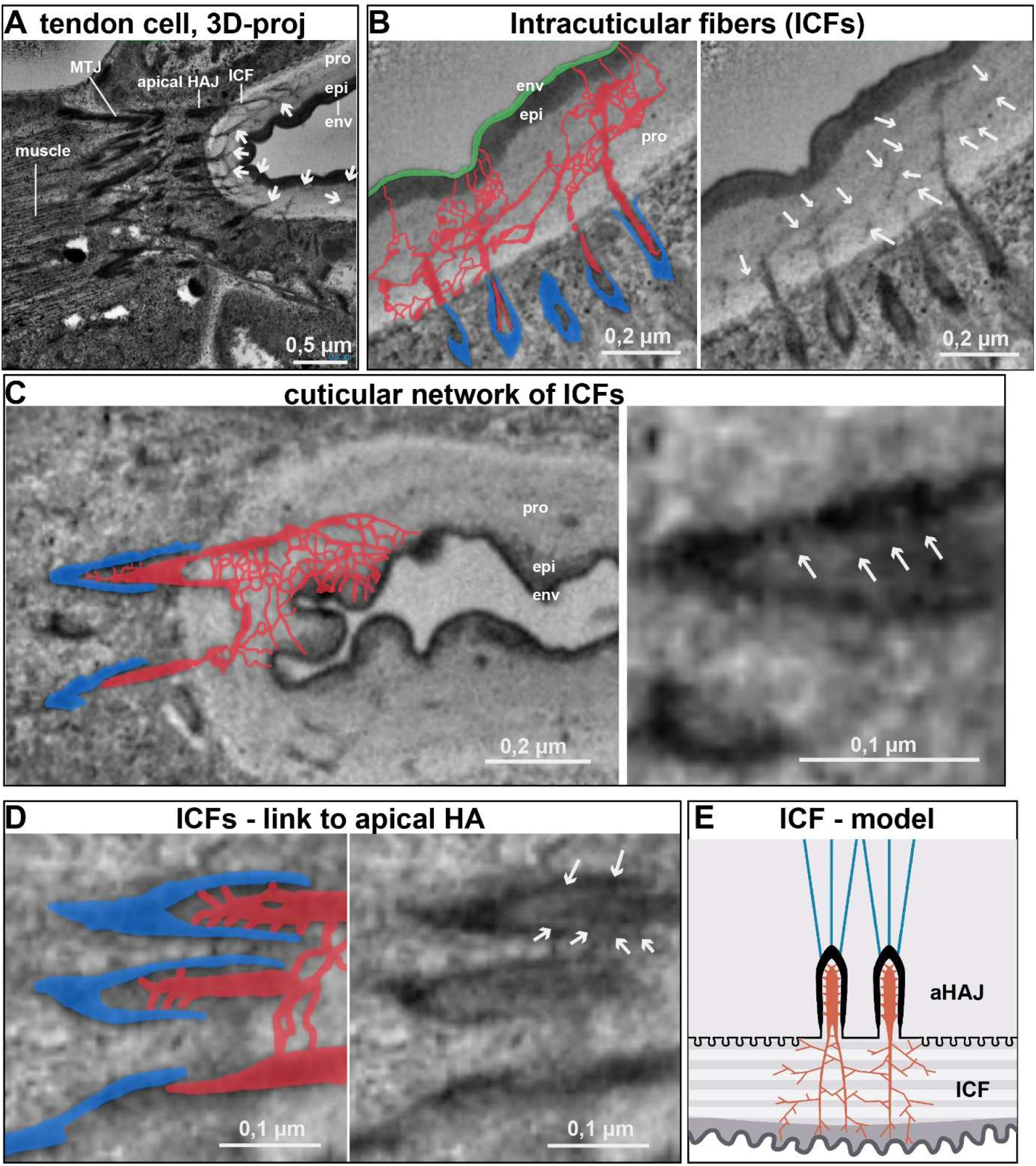
Intracuticular fibers of the musculature and their attachment sites to the epidermal cuticle. **(A)** 3D projection reveals branching of ICFs within the cuticle (posterior segment) and their contact points at the epidermal tendon cell and muscle. White arrows point to the ICFs. Abbreviations: apical Hemi-Adherens Junctions (aHAJs), env (envelope), epi (epicuticle), ICFs (intracuticular fibers; tonofilaments), myotendinous junctions (MTJs), pro (procuticle). **(B, C)** Adjacent tonofilaments are labeled: red for tonofilaments, blue for aHAJs, and green for the envelope. Branching of ICFs and (f)-ICFs in the procuticle is visible. Unlabeled images of ICFs and f-ICFs are provided in Fig. S8. Arrows in the right image point to filamentous ICFs in the cuticle (B) or connected to the aHAJ membrane (C). **(D)** Magnified view of aHAJs from C shows multiple f-ICF attachments to the aHAJ membrane. The right image is unlabeled. **(E)** The model indicates the pattern of ICFs (orange) in the tendon cell cuticle; microtubuli of tendon cells are marked in blue. Scale bars are indicated.

### Reconstruction of larval intracuticular fibers (ICFs) at chordotonal organs

Chordotonal organs stretch subepidermally to detect mechanical stimuli. Ligament and cap attachment cells connect these organs to the epidermal cuticle, which is evident in 2D TEM images and 3D reconstructions (Fig. 10A; Fig. S8A-C). Thus, ligament and cap cells experience considerable mechanical stress, which impacts their attachment to the cuticle. To better understand their attachment site at the cuticle, we analyzed 3D projections of the chordotonal organ cuticle. We observed electron-dense filamentous structures that reach from the attachment cells into the cuticle (Fig. 10B; Fig. S8D, Fig. S9). These filamentous structures resembled the structure of ICFs of tendon cells. Therefore, we will refer to them as chordotonal organ ICFs (intracuticular fibers). Similar to tendon cells, we found large ICFs and filamentous ICFs in the chotodotonal attachment cell cuticle. Large ICFs are anchored via numerous filamentous ICFs to aHAJs-like structures at the apical cell membrane (Fig. 10C). Within the cuticle numerous filamentous-like ICFs branch from the large ICFs in a network-like pattern (Fig. 10D). All ICFs likely form an anchorage throughout the cuticle layers. These intricate mechanical assemblies (Fig. 10E) closely mirror the architecture found in tendon cells at muscle attachment points (Fig. 9E). Furthermore, the cuticle attachment cells exhibit arrays of parallel microtubule bundles that extend between basal cell membrane junctions and the apical aHAJs, where the ICFs anchor (Fig. S9). Overall, our data indicate that the ICFs of tendon cells and the chordotonal organs-ICFs are densely anchored in the cuticle and at the apical cell membrane, likely to withstand mechanical stress and possibly to sense it.

**Figure 10.**
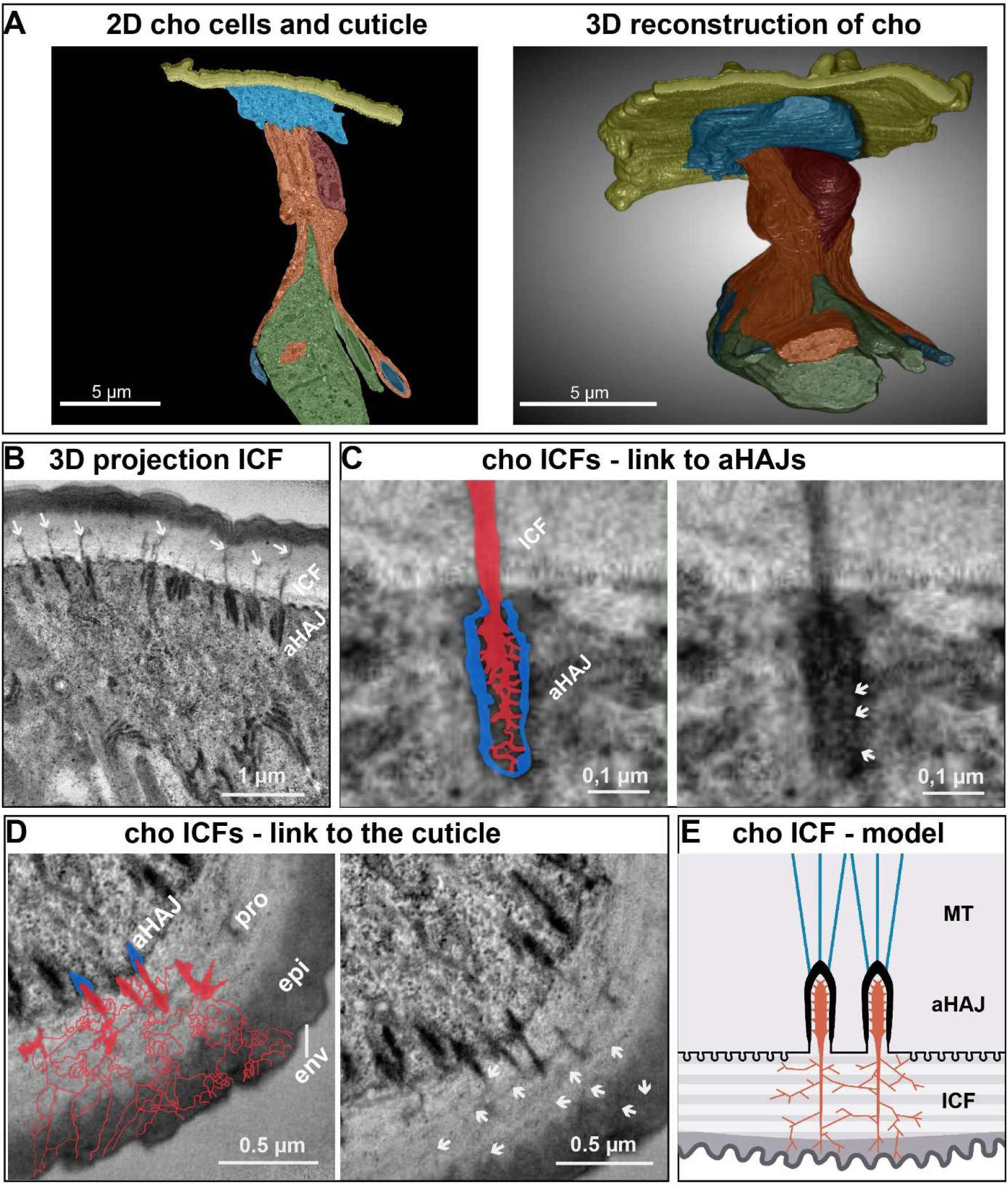
The chordotonal organ (cho) and its attachment site on the epidermal cuticle. **(B)** Cells of chordotonal organ and a 3D reconsctruction, which shows the ligament attachment cell (blue) firmly attached to the cuticle over a large area. Extending from it, the glia (sheath) cells (brown) are visible, and finally, the axon (green) and three dendrites branching off from it. **(B-D)** Chordotonal organ attachment cells contain thick intracuticular fibers (ICF, marked with red arrows in B) and thinner filamentous ICFs (marked in red and with arrows in D). Numerous ICFs extend into the porcuticle (pro), epicuticle (epi), and envelope (env) layers. At the cell membrane ICFs are connected (arrows in C) to aHAJs (marked in blue in C). Other images and 3D projections of the ICF are provided in Figs. S9 and S10. **(E)** The model shows the pattern of chordotonal organs ICFs (orange) in the attachment cell cuticle. Microtubuli (MT) are marked in blue. Scale bars are indicated.

## Discussion

Despite advances in microscopic and chemical imaging techniques uncovering cuticle-specific formation and functions ^3,10,17,22,28,38,40,67–69^, the fine anisotropic structures in the insect cuticle’s three-dimensional architecture remain poorly understood. We combined the serial TEM dataset with deep learning-based 3D modeling to explore the intricate micro- and nanostructures of the aECM at different insect epithelial organs. The established dragonfly convolutional Neural Network (CNN) excels at detecting subtle, microstructural anomalies and features ^70–74^, thriving on the natural complexity of patterns that traditional machine learning often falters on. Harnessing cutting-edge AI-driven computer modeling, we analyzed a vast dataset to reconstruct the complex architecture and surface textures of the cuticle in detail. This approach unlocks precise 3D analysis with spatial accuracy, transforming abstract and nanoscale structures into vivid, visually accessible forms and saving significant time through semi-automated analysis (Figs. 1; Figs. S1,S2). Our results highlight the 3D reconstruction of key morphological features of the exoskeletal cuticles. By generating detailed 3D models of the epidermal cuticle (Fig. 1,2), sensory organs (Fig. 3), tracheal system (Fig. 4), spiracles (Fig. 5) in the *Drosophila* first instar larva, along with ultrastructural 3D projections of cellular cuticle adherens in muscles and sensory organs (Figs. 9,10) and the characterization of epidermal and anal pad cuticular pore canal-like systems (Figs. 6-8), we provide a foundation for future genetic, evolutionary, and bioengineering research.

Previously, Richter et al. used this data volume to establish a proper classification of sensilla based on external and internal morphology. The larval body wall contains 541 sensory cells that are linked to external sensilla. Of these, 363 are likely mechanosensory, identified by their tubular bodies. These mechanosensory cells are distributed among 201 papilla sensilla, 100 hair sensilla, and 22 knob sensilla, with an additional 40 located in the major head organs and the spiracle sense organ ^18^. In contrast, we focused on the 3D surface features of the cuticle at the sensilla. As expected, we found that the anterior region of the larva’s epidermal head contains thermosreceptors, mechanoreceptors and chemoreceptors, which are essential for detecting environmental stimuli and orienting the larva (Fig. 2). We identified cirri, composed of fine sensory bristles near the mouth, which support tactile perception and help the larva locate and guide food toward the mouth. We analyzed the main types of sensilla most common on the larval body wall: papilla, hair, and knob sensilla. These sensilla appear either individually or as part of sensory organs or compound sensilla such as the terminal organ, Keilin’s organ, or terminal sensory cones (Fig. 2). Our studies on the reconstruction of the epidermal cuticle confirm that the chitin matrix in the sensory organs determines the pathway for stimulus transmission. Ultrastructural examinations of longitudinal sections along the epidermis show that sensilla extend deep into the chitinous porcuticle, and in the case of papilla and knob sensilla, reaching even the epicuticle. At the hair sensilla, the cuticle appeared even thicker than in the adjacent epidermal cell regions (Fig 3). This likely reflects the sequence of cuticle secretion and sensilla growth, as in embryos, the outermost envelope and epicuticle form first, followed by the chitin-based procuticle ^75^. During cuticle secretion, the apical regions of provisional accessory cells transform to form the cuticular apparatus, which receives sensory stimuli ^63^. Thus, we characterized the morphological structures that demonstrate the close intertwining of sensory and mechanical elements in larval development. This dataset provides a basis for understanding the larva’s functional organization and its interactions with the environment

The cuticle is permeable to oil-soluble molecules, including pheromones, telergones, hormones, and insecticides ^76^. Rich in proteins and lipids, the cuticle’s outer layers play an essential role in its physical and chemical properties ^24^, providing a barrier to insecticides and chemical penetration ^77^. This prompts the question of whether and how underlying cells regulate these layers through the large chitin matrix. Recent studies confirm that these channels are essential for the transport of proteins and lipids^27,40^. Deformation of the pore canals can reduce cuticle lipids, thereby increasing permeability to water and xenobiotics, indicating a compromised barrier function ^27,40^. Pore canals can contain numerous thin wax canal-like structures, which emerge from the pore canals, pass through the epicuticle, and reach the outer layer to deliver lipids/waxes ^78,79^. These canals also penetrate laterally between the lamellae of the procuticle ^79,80^. Our 3D reconstructions characterize potential pore canal-like structures in the *Drosophila* larva and confirm that these nano-canal structures extend from the membrane into the chitinous procuticle with external diameters below 20 nm (Fig. 7), which is consistent with previous ultrastructural analyses of other insects. Further, our reconstructions reveal that these structures occur in large numbers, likely establishing a network-like organization across the cuticle layers in the *Drosophila* larva (Fig. 7,8). Such a network of pore canals suggests that the cuticle may undergo a dynamic process that transports chitin, lipids, and proteins into the different layers and across to the surface for diverse dynamic subcellular processes.

Recent FIB-SEM imaging studies reveal that multiple channels extend from the apical cell membrane ^43^. However, so far, studies have focused on insects with large pore canals, whereas those in *Drosophila* larvae are thin and difficult to identify ^23^. The whole-animal ultrastructural volume enabled us to characterize the overall organization of these critical pore and wax canals in the epidermal cuticle. In the 3D projections, we observed thicker pore canal-like structures in the chitin matrix from which thin structures extend into all cuticle layers. Our research indicates that *Drosophila* pore canals not only cross-link but also weave an intricate network of fine, thin canals throughout the cuticle. This elaborate network may take shape within the chitin matrix and may stretch from the epicuticle to the outer envelope (Fig. 7). Such a labyrinth of thin canals might not only be a transport system that acts as a conduit for proteins, lipids, and polyphenols ^79,81^, shuttling them to their destinations, but perhaps also keep fluids and lipids that help the cuticle adapt to metabolic shifts or mechanical challenges.

The anal pads primarily function as ion transport organs, actively taking up ions from the surrounding environment into the larval body. Larval anal pad cells are highly specialized, characterized by a thin cuticle, enlarged membrane infoldings linked to ion transport. Their highly differentiated, transport-active cells feature large, sheet-like infoldings of the plasma membrane beneath the cuticle ^42^. The cuticle of these cells features epicuticular depressions, rather than the typical pore-canal system found in the epidermis ^82,83^. Thus, a thinner cuticle might be adapted for Ion-transport and osmoregulation, with little emphasis on pore canals. We observed only the envelope adjacent to the chitinous procuticle in the anal pad, but no epicuticle. Moreover, we observed large, electron-dense, bag-like structures extending from the envelope into the chitin matrix, with multiple potential thin canals branching toward the apical cell membrane (Fig. 6; S5). These bag-like structures may reflect cylindrical tubules in the dense layer of the epicuticle of Musca domestica larvae ^83^. Since pore canals are responsible for transport through the cuticle, it is tempting to speculate that their specific structural features may support the transport activity of the anal pads. In contrast, epidermal cells primarily transport proteins and lipids to the outer layers and possibly molting fluid through the canals, while also preventing fluid loss and thus dehydration; epidermal canals may differ in nanostructure from those of the anal pads.

Serving as the first line of defense against external influences and mechanical stress, the cuticle must not only be water-repellent and selectively permeable, but also firmly anchored in the underlying cell layer. Cuticle adhesion to the cells occurs via the connection of the apical hemiadherens junctions on the apical cell membrane, from which electron-dense cuticular fibers extend into the procuticle. Genetic knockdown and mutant studies demonstrated that defects in chitin matrix organization and insufficient chitin deacetylation weaken this adhesion ^17,54^. The effects are even worse when the ZP matrix in the cuticle is lost, leading to the loss of the ICF and detachment of the cuticle ^17,21^. In all cases, the animals’ mobility is severely restricted because ICFs connect the cuticle primarily to tendon cells at the apical membrane, while muscles attach to the basal side. As a result, ICFs can experience significant mechanical stress during normal locomotion. Until now, the mechanism by which ICFs are anchored in the cuticle remained unclear. Our 3D analyses identify a dense network of thread-like filaments ICFs extending from large ICFs reaching into all cuticle layer, forming a robust structure that may secures the system within the cuticle (Fig. 9). Our data further suggest that mechanoreceptors like the chordotonal organs, stretching across the epidermis much like somatic muscles, are secured to the cuticle through a similar anchoring system (Fig. 10). Ligament and cap attachment cells together form a robust internal cytoskeleton that ends at aHAJs on the apical cell membrane from where large IFCs extend into the chitin-containing procuticle, which are connected through a web of filamentous ICFs to the cuticle layers (Fig. 10). To sum up, the cuticle is anchored to tendon cells or chordotonal organs through a dense, protein-packed network that adheres to the apical cell membrane, is reinforced by ICFs and f-ICFs, and relies on a sturdy internal cytoskeleton for support.

Nature creates diverse material properties, such as flexibility, stiffness, and toughness, by organizing components across multiple hierarchical levels rather than relying solely on chemical composition ^15^. Biological materials use a limited set of biopolymers, minerals, ions, and cross-linking agents ^75^. The mechanical properties of cuticles, including modulus (E), depend on moisture content, chitin or chitosan content, protein crosslink density, lipids, and mineral particles ^7,24,85^. Chitin, the primary component of the cuticle, is biodegradable and biocompatible, making it suitable for even medical applications. However, chitin crystals are inherently rigid and brittle. In contrast, the insect cuticle demonstrates flexibility, stiffness, strength, and bioactivity. Insects achieve these properties by linking chitin fibrils with proteins, enzymatically trimming them, and partially converting chitin to chitosan through deacetylation. What gives the cuticle its remarkable ability to endure intense mechanical forces in these organs? One key lies in the chitin matrix, where fiber orientation dictates both the direction and the degree of its strength. When these chitin fibers run parallel to the direction of tension, the cuticle becomes stronger along that line than across them ^75^. The Bouligand architecture of chitin fibers maximizes energy dissipation by promoting crack bridging, deflection, arrest, and twisting ^76,77^. Yet, our research suggests that neither chitin-fiber orientation nor tensile strength alone fully covers the cuticle’s remarkable stability. The massive pore canal tubules, which we observed in the helicoidal structure of the epidermal aECM, can serve an important mechanical role by imparting anisotropy and toughness ^78–80^. Another key lies in a sophisticated anchoring network, in which coarse and fine fibers bind the cuticle securely to its underlying substrate. This union of a soft cell membrane with a resilient chitin matrix transforms the cuticle into a functional exoskeleton. When this anchoring fails, the chitin matrix separates, impairing the animal’s usual mobility ^23^. Our findings suggest that biomimetic materials with diverse properties could be anchored just as effectively by emulating this natural fiber system.

## Material and Methods

### Whole Volume data

All reconstructions were done on a STEM (transmission electron microscopy) volume of a whole first instar larva; information on the technical details of its generation is published _recently_ 18,56,57

### Image Processing

Image processing was performed using HP Z4 workstations equipped with Intel Xeon processors and Nvidia GeForce GTX card (16GB). A primary system with 128 GB RAM was used for standard processing tasks, while a second workstation with 256 GB RAM was employed for computationally intensive operations to ensure efficient handling of large-scale datasets.

The dataset consisted of 4816 JPEG images of a first-instar larva of *Drosophila melanogaster*, acquired using transmission electron microscopy (TEM). The total dataset size was approximately 3.5 terabytes, with each image having a resolution of 99600 × 31600 pixels. The dataset was provided by the Thum lab ^18^. This subdivision was maintained throughout the study due to memory and computational limitations.

At the beginning of the study, attempts were made to open and process the image files using ImageJ and Imaris 10.01; however, these efforts were unsuccessful due to performance limitations and constraints in data import, merging, and segmentation. Dragonfly Pro was used for its efficient AI-based processing of electron microscopy data. Image conversion and resizing were performed using XnConvert (version 1.100.1). Subsequent image analysis, segmentation, and 3D visualization were performed in Dragonfly Pro, while final image editing was conducted in Adobe Photoshop CS6.

For processing, the dataset was divided into three anatomical regions: posterior and medial (each 40000 × 31600 pixels), and anterior (19600 × 31600 pixels). To enable processing in Dragonfly Pro, the images were resized by a factor of 10 using XnConvert. The posterior and medial datasets were reduced to 4000 × 3160 pixels, while the anterior dataset was reduced to 1960 × 3160 pixels, preserving the highest possible resolution.

To isolate specific cuticular structures, such as hair sensilla, high-resolution regions of interest (ROIs) were extracted using the cropping tool in XnConvert. These regions were subsequently used for detailed 3D reconstruction.

The processed image datasets were imported into Dragonfly Pro and handled in three separate blocks (posterior, medial, and anterior; 4816 images each). All datasets were calibrated with a voxel size of x = 0.05 µm, y = 0.05 µm, and z = 0.035 µm. Due to hardware limitations, a high-performance workstation (256 GB RAM) was required for efficient data processing.

Following import and calibration, 3D stitching of the datasets was attempted. However, stitching all three datasets simultaneously exceeded the GPU’s capabilities, leading to processing errors and software instability. Therefore, only the medial and anterior datasets were successfully stitched, while the posterior dataset was appended without stitching. Subsequent analyses were conducted on two datasets (anterior–medial and posterior), thereby circumventing hardware limitations.

### Volume Segmentation

Segmentation of the epidermal and tracheal cuticle was performed using a deep learning– based approach. Manual segmentation of all slices was not feasible due to the dataset size. Initial attempts using the Segmentation Wizard and conventional machine learning methods yielded unsatisfactory results in terms of accuracy and reproducibility. Therefore, eleven representative slices from different regions of the dataset (beginning, middle, and end) were selected as training data. Multi-region-of-interest (ROI) annotations (cuticle, trachea, and background) were manually generated using the painter tool. Based on these annotations, a deep learning model was trained in Dragonfly Pro to enable automated segmentation.

For high-resolution reconstruction of specific structures, including single and double hair sensilla, knob and papilla sensilla, posterior spiracles, and the spiracular chamber with Filzkörper, a separate deep learning model was trained using the same procedure.

Post-processing was performed using the smoothing tool in Dragonfly Pro to eliminate segmentation artifacts and misclassifications, which were primarily caused by structural similarities between imaging artifacts and biological features. This step significantly improved segmentation accuracy and continuity. Finally, segmented datasets were rendered into 3D models. Visual optimization was achieved through the application of lighting and shading techniques to enhance structural clarity and overall representation.

### FIB-SEM Imaging

Adult wildtype *Drosophila melanogaster* larvae were placed in a crossing cage for embryo collection on pre-prepared apple agar embryo collection petri-dishes. After 2 hours of egg-laying 20-30 embryos are taken away from the crossing cages and allowed to hatch in 25°C and 60% relative humidity. The next day (22-24 hours) 10 first instar larvae are dissected in two ways; five 1^st^ instar larvae are dissected at the anterior end by cutting the head, and five are dissected at the posterior end by cutting the anus. Each dissected specimen is then fixed in 2% paraformaldehyde (PFA; EMS #19208) and 2% glutaraldehyde (GA; EMS #E16220) in 100 mM phosphate buffer (PB) for 2 hours at room temperature (RT). Samples then washed three times for 5 min each in 100 mM HEPES (Sigma-Aldrich #H3357), followed by three washes for 5 min each in pure water (p.H₂O).

Heavy-metal staining proceeded as follows: First osmication: 30 min at RT in a 1:1 mixture of 3% potassium hexacyanoferrate (VWR #26816.232) in 3 mM CaCl₂-dihydrate (Merck #102382) and 4% OsO₄ (EMS #E19190), yielding a final volume of 1 mL (0.015 g potassium hexacyanoferrate in 500 µL of 4 mM CaCl₂ + 4% OsO₄). Washed 3 × 5 min at RT with p.H₂O. Thiocarbohydrazide (TCH): 0.1 g TCH (Sigma-Aldrich #223220-5G) dissolved in 10 mL p.H₂O, submerged for 20 min at RT, then incubated for 1 hour at 60 °C with 10 min stirring, followed by filtration (Carl Roth CHROMAFIL PET syringe filter #P298.1). Washed 5 × 5 min at RT with p.H₂O. Second osmication: 30 min with 2% OsO₄. Washed 3 × 5 min at RT with p.H₂O. Uranyl acetate: Overnight incubation at 4 °C in 1% uranyl acetate (EMS #E22400). Next day: washed 3 × 5 min at RT with p.H₂O. Dehydration and embedding. Samples were dehydrated in graded ethanol/acetone series (Ethanol absolut ≥99.8%, AnalaR NORMAPUR® ACS #20821.296; Acetone, EMS #10016) at concentrations of 30, 50, 70, 90, 96%, and 3 × 100% EtOH (20–30 min per step). Infiltration with Epon (EMS EPON-Araldite #E70900) was performed over 2 hours in decreasing ethanol concentrations (25, 50, 75% EtOH), followed by overnight incubation in pure Epon at RT. Final embedding lasted 4–6 hours in pure Epon and cured for 48 hours at 60 °C in silicone molds (Easy-Molds™ #E69931-05). Mounting and coating. Cured blocks were trimmed and polished using an ultramicrotome (Leica UC6, Wetzlar, Germany), then cut to size and mounted on aluminum stubs (Micro to Nano #10-0030120100) secured with silver-based conductive paint (PLANO GmbH #D35578) on double-faced carbon tape (Micro to Nano #15-000412). Samples were sputter-coated with 80/20 platinum/palladium for 20 seconds at 40 µA using a Q150T ES sputter coater (Quorum Technologies, Lewes, UK). Samples were stored in a vacuum chamber and placed inside the FIB-SEM at least 12 hours before imaging.

Serial-section imaging was performed on a Crossbeam 550 (Zeiss, Germany). SEM imaging was conducted at 4 kV acceleration voltage and 200 pA imaging probe current using backscattered electron (ESB) and InLens secondary electron detectors simultaneously. FIB milling was performed at 30 kV accelerating voltage with a 700 pA probe current. Automated serial imaging was controlled using Atlas 3D software (Fibics Incorporated, Ottawa, Canada). Each scan have a voxel size of (5 × 5 × 10) nm.

## Acknowledgemets

We gratefully acknowledge Politi’s Lab at TU Dresden for providing access to the equipment essential for conducting the FIB-SEM experiment. We thank Dr. Thomas Kurth and Susanne Kretschmar from the Core Facility Electron Microscopy and Histology of the Center for Molecular and Cellular Bioengineering Technology Platform (CMCB TP, University of Technology Dresden) for FIB-SEM sample prep. We thank Thum’s and Behr’s labs for technical support and discussions on the data and manuscript. MB thanks the Deutsche Forschungsgemeinschaft (DFG) for support (BE3215/5-1).

